# Anhedonia following mild traumatic brain injury in rats: A behavioral economic analysis of positive and negative reinforcement

**DOI:** 10.1101/527895

**Authors:** Pelin Avcu, Ashley M. Fortress, Jennifer E. Fragale, Kevin M. Spiegler, Kevin C.H. Pang

**Author notes:** Authors contributed equally (Pelin Avcu, Ashley M. Fortress, Jennifer E. Fragale, Kevin M. Spiegler). VA Pittsburgh Healthcare System, University Drive C, Building 30, Pittsburgh, PA 15240. Rutgers University/Rutgers Biomedical and Health Sciences, 683 Hoes Lane West, Office 259A, Piscataway, NJ 08854. Corresponding Author: Kevin C.H. Pang, Ph.D., New Jersey Health Care System, 385 Tremont Avenue, Mailstop 15, East Orange, NJ, 07018.

## Abstract

Psychiatric disorders affect nearly 50% of individuals who have experienced a traumatic brain injury (TBI). Anhedonia is a major symptom of numerous psychiatric disorders and is a diagnostic criterion for depression. It has recently been appreciated that reinforcement may be separated into consummatory (hedonic), motivational and decisional components, all of which may be affected differently in disease. Although anhedonia is typically assessed using positive reinforcement, the importance of stress in psychopathology suggests the study of negative reinforcement (removal or avoidance of aversive events) may be equally important. The present study investigated positive and negative reinforcement following a rat model of mild TBI (mTBI) using lateral fluid percussion. Hedonic value and motivation for reinforcement was determined by behavioral economic analyses. Following mTBI, the hedonic value of avoiding foot shock was reduced. In contrast, the hedonic value of escaping foot shock or obtaining a sucrose pellet was not altered by mTBI. Moreover, motivation to avoid or escape foot shock or to acquire sucrose was not altered by mTBI. Our results suggest that individuals experiencing mTBI find avoidance of aversive events less reinforcing, and therefore are less apt to utilize proactive control of stress.

## 1. Introduction

Traumatic brain injury (TBI) is one of the leading causes of disabilities and deaths worldwide and in the US [1]. Among the disabilities, psychiatric disorders are common, estimated to occur in nearly 50% of those experiencing a TBI [2]. Moreover, major depression is observed in approximately 40% of those hospitalized for TBI [3]. Thus, understanding the mechanisms by which TBI leads to depression and other psychiatric disorders is increasingly necessary.

Anhedonia is a prominent feature of depression and may be important in the co-morbidity that is often observed with anxiety disorders [4]. Recently, the study of reinforcement has indicated multiple components of reinforcement processing [5, 6]. Consummatory (or hedonic), motivational and decisional components of reinforcement processing can be independently altered in psychopathology [7]. For this manuscript, the term anhedonia is used to refer to decreases in hedonic value (consummatory component); motivational and decisional components refer to motivation and decision-making, respectively. Patients with schizophrenia have relatively normal hedonic experiences (consummatory component), but are impaired in making optimal decisions when comparing values of different items (decisional component) [8]. Addiction is associated with increased desire (motivational component) and activation of motivational brain circuits in response to drug cues [9]. Finally, individuals with depression have reduced motivation to work for monetary reward [10]. The identification of the components of reinforcement processing that are disrupted by TBI may allow for better treatment of TBI-induced depression.

The study of reinforcement processing in experimental models of TBI are relatively few. Controlled cortical impact (CCI) injury to the medial frontal cortex reduced the preference for saccharine 8 days after injury, and CCI injury of the parietal cortex reduced sucrose preference 5.5 months after injury [11, 12]. Similarly, lateral fluid percussion injury reduced sucrose preference 4 days after injury, an effect that was reversed by fluoxetine. Midline fluid percussion reduced preference for sucrose 30 days after injury when combined with an immune challenge of lipopolysaccharide (LPS) administration, as compared to an LPS sham group [13, 14]. In contrast, other studies found that lateral fluid percussion injury did not alter sucrose preference at 1, 3 or 6 months after injury, and neither mild nor severe CCI altered preference for sucrose 16 days after injury in mice [15, 16]. Unfortunately, the use of the sucrose or saccharine preference test does not assess different components of reinforcement. Therefore, the present study utilized behavioral economic analysis to investigate changes in consummatory and motivational components of reinforcement following mTBI [7, 17].

Behavioral economics is based on the premise that consumption varies as the price of goods changes with the idea that consumption decreases as price increases [18]. Behavioral economic analysis allows for the separate assessment of hedonic and motivational components of reinforcement [19]. Consumption (or demand) at a theoretical price of 0 is represented by *Q*_0_ and is related to the hedonic value of the reinforcer. Demand elasticity denoted by _α_ describes the relationship between consumption and price and is inversely related to motivation. Behavioral economic analysis has been widely used to interpret these components for a wide variety of positive reinforcement [17, 19–22].

Reinforcement can be divided into positive and negative reinforcement. Obtaining a desirable stimulus (positive reinforcement) or removal of an undesirable stimulus (negative reinforcement) increases the likelihood of repeating the response that produced the reinforcement. With regard to psychopathology, much more is known about positive reinforcement compared to negative reinforcement. Positively-valenced items have lower ratings and are less arousing to individuals with depression [7]. Whether the symptom of anhedonia in depression is specific to positively-valenced items or due to a global flattening of both positive and negative valence items needs further investigation [23]. Similarly, anxiety is often associated with negative bias, whereby individuals have enhanced attention to and motivation to remove negatively-valenced items [24–27]. Importantly, anhedonia was a predictor for time to remission, was inversely correlated with depression-free days in SSRI-resistant adolescents, and may be a predictor of successful outcome following intravenous subanesthetic ketamine treatment in patients with treatment-resistant depression [28, 29]. Therefore, the study of reinforcement may prove insightful in understanding mood disorders caused by TBI.

Stress often results from encountering aversive events and can be a precipitating factor in depression and anxiety [30, 31]. Cognitive control refers to the adaptive changes required to accomplish our goals in the face of challenges and conflicts, and impacts emotion and stress regulation [32]. Cognitive control includes both reactions to current stressful events (reactive control) and in anticipation of a future stressful event (proactive control), each of which may use different but potentially overlapping brain regions [33]. Critically, reduced proactive control may limit individuals with depression to successfully cope with stress [34].

Recently, we applied behavioral economic analysis to negative reinforcement where animals could escape foot shock [35] or avoid foot shock [36]. The hedonic value of shock termination (*Q*_0_, “relief”) increased as the intensity of the shock increased, and motivation to escape shock increased as shock intensity increased (_α_ decreased). In addition, a rat model of anxiety vulnerability (Wistar Kyoto rat) was more motivated to escape or avoid foot shock compared to the outbred Sprague Dawley rat. The results demonstrate that negative reinforcement can be separated into hedonic value and motivation using behavioral economics, similar to that done for positive reinforcement. Furthermore, escape and avoidance of foot shock have parallels to reactive and proactive control of stress, respectively.

The present study evaluated the effects of mTBI on positive and negative reinforcement using behavioral economic analyses. Based on prior studies that investigated the effects of TBI on sucrose preference, it was hypothesized that mTBI would cause anhedonia or decreased motivation for positive reinforcement. No prior studies have utilized behavioral economic analysis to separate different components of reinforcement processing following mTBI. Because of the lack of studies in this area, the present study will provide novel information on the aspects of positive and negative reinforcement that are impaired following mTBI with potential relevance to treatment of depression.

## 2. Materials and Methods

### 2.1. Animals

Forty-four male Sprague Dawley rats (approximately 3 months of age, 250 – 300 g at the start of the studies; Envigo, Indianapolis, IN) were housed individually in a room with a 12:12 hour light:dark cycle. One rat did not learn to lever press for sucrose and was removed from the study, resulting in 6 Sham and 5 TBI rats in Experiment 1. Two rats died after the TBI/Sham surgery, resulting in 7 Sham and 8 TBI rats for Experiment 2 and the same numbers for Experiment 3. Food and water were available *ad libitum*, except during training and testing with sucrose reinforcement. For procedures using sucrose reinforcement, rats were restricted of food to maintain a body weight that was 85% of the *ad libitum* weight. All procedures were conducted in accordance with the NIH Guide for the Care and Use of Laboratory Animals and approved by the IACUC of the Veterans Affairs Medical Center at East Orange, New Jersey.

### 2.2. Acoustic Startle Reflex

All rats were assessed for acoustic startle reflex (ASR) because we previously found suppression of ASR to be a consistent outcome of mTBI following lateral fluid percussion injury [37–39]. Assessment of ASR was performed as previously described [37]. ASR was assessed in individual, sound-attenuated chambers, each equipped with an exhaust fan, a speaker to maintain background noise (68dB) and another speaker for delivery of the acoustic stimuli (86, 96, and 102dB; 100 ms). A single session consisted of 24 trials (8 repetitions of 3 intensities), in which the acoustic stimulus was presented immediately after a 250ms baseline period. Trials occurred every 25-35 seconds in a pseudo-randomized order. Rats were loosely restrained on platforms that transduced whole body force (Coulbourn Instruments, Holliston, MA). Movement was considered a startle response if the magnitude exceeded the sum of the maximum magnitude during the baseline period and 4x the standard deviation of baseline activity. Two dependent measures were obtained: sensitivity and magnitude. ASR sensitivity was the probability of a startle response to an acoustic stimulus (expressed as a percentage at each stimulus intensity). ASR magnitude was the amplitude of the startle response at each stimulus intensity, and was corrected for the subject’s body weight. ASR magnitude was calculated only for those trials in which a startle response was detected. If no responses were elicited for all trials of a stimulus intensity, ASR magnitude was assigned a value of 0 for that subject. ASR was measured before injury, and 1, 7 and 14 days after injury. The pre-injury startle magnitude at 102 dB was used to match rats and then randomly assign matched rats to sham and mTBI groups.

### 2.3. Surgery

All rats underwent TBI/sham procedures prior to behavioral training and testing (except for ASR). Rats were prepared for lateral fluid percussion injury, as previously described [37, 40]. Briefly, rats were anesthetized with a mixture of ketamine and xylazine (60 mg/7 mg per kg body weight, 1 ml/kg, i.p.). A craniectomy was created in the left or right parietal bone plate (4 mm diameter, centered at −3.0mm posterior to and ±3.5mm lateral to Bregma). Left and right craniectomy locations were counterbalanced across animals. A Luer-Lok connector was glued to the skull surrounding the craniectomy. A plastic cylinder (cut from a 12 ml syringe) surrounded the craniectomy to protect the Luer-Lok connector. Dental cement was used to stabilize the plastic cylinder and Luer-Lok connector to the skull. A small piece of Kimwipe was inserted into the connector to keep the dura clean of debris.

### 2.4. Fluid Percussion Injury

mTBI was produced using lateral fluid percussion, a well-established experimental model of TBI that produces a focal injury, modeling a clinical contusion injury without skull fracture [41–43]. Subjects were anesthetized with 5% isoflurane and Luer-Lok hub connected to the device. As soon as rats recovered a foot-pinch reflex bilaterally, a fluid pressure pulse was delivered to the dura using a computer controlled voice-coil injury device (Table 1) [44]. After delivery of the impact, duration of apnea was recorded, rats were laid supine and the latency of their righting reflex (in the absence of any tactile stimulation) was recorded. Sham control subjects also underwent surgery and isoflurane anesthesia, and were attached to the device. However, a fluid pressure pulse was not delivered. Therefore, only latency of the righting reflex was recorded for Sham controls as a measure of the effects of anesthesia alone.

**Table 1.**
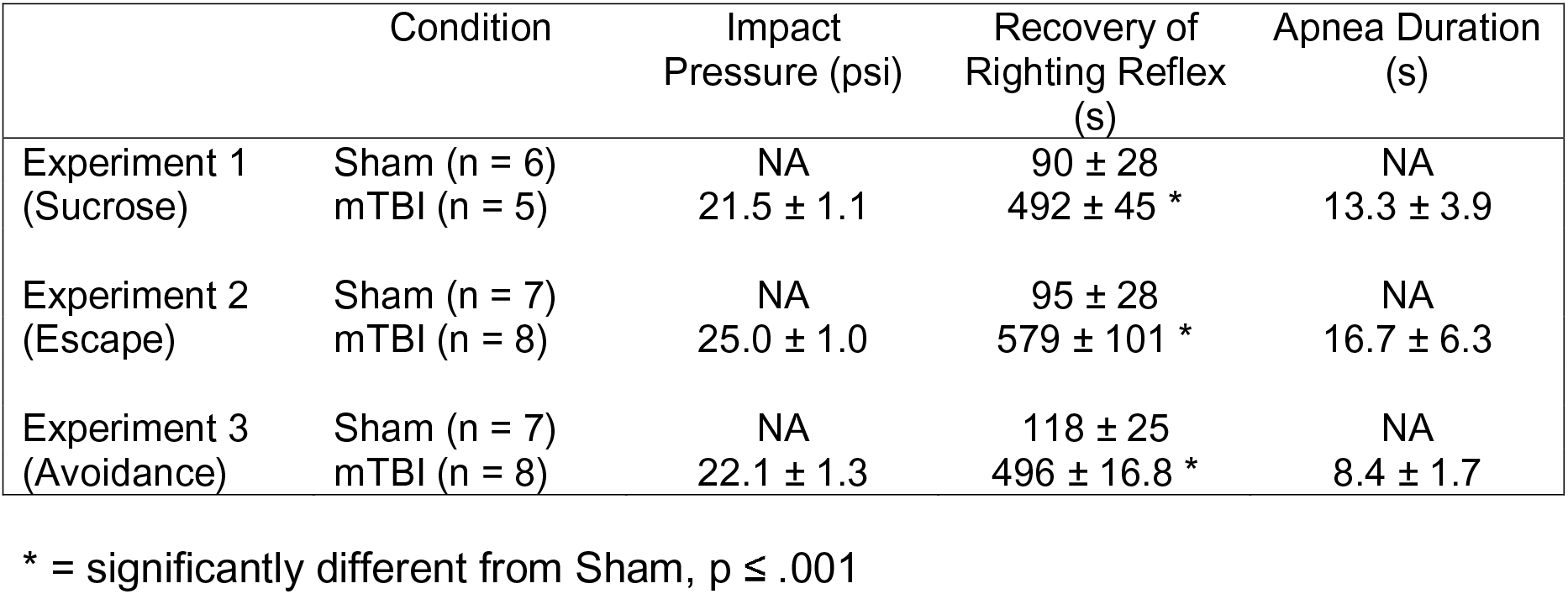
Injury pressure and acute signs. Mean ± S.E.M.

### 2.5. Behavioral Training and Testing

The primary focus of these studies was to assess demand characteristics of positive and negative reinforcement after mTBI. We were interested in the post-acute phase so the demand curve was obtained between 3 weeks to 1 month after injury. However, additional testing also occurred prior to this time, as detailed in Table 2.

**Table 2.**
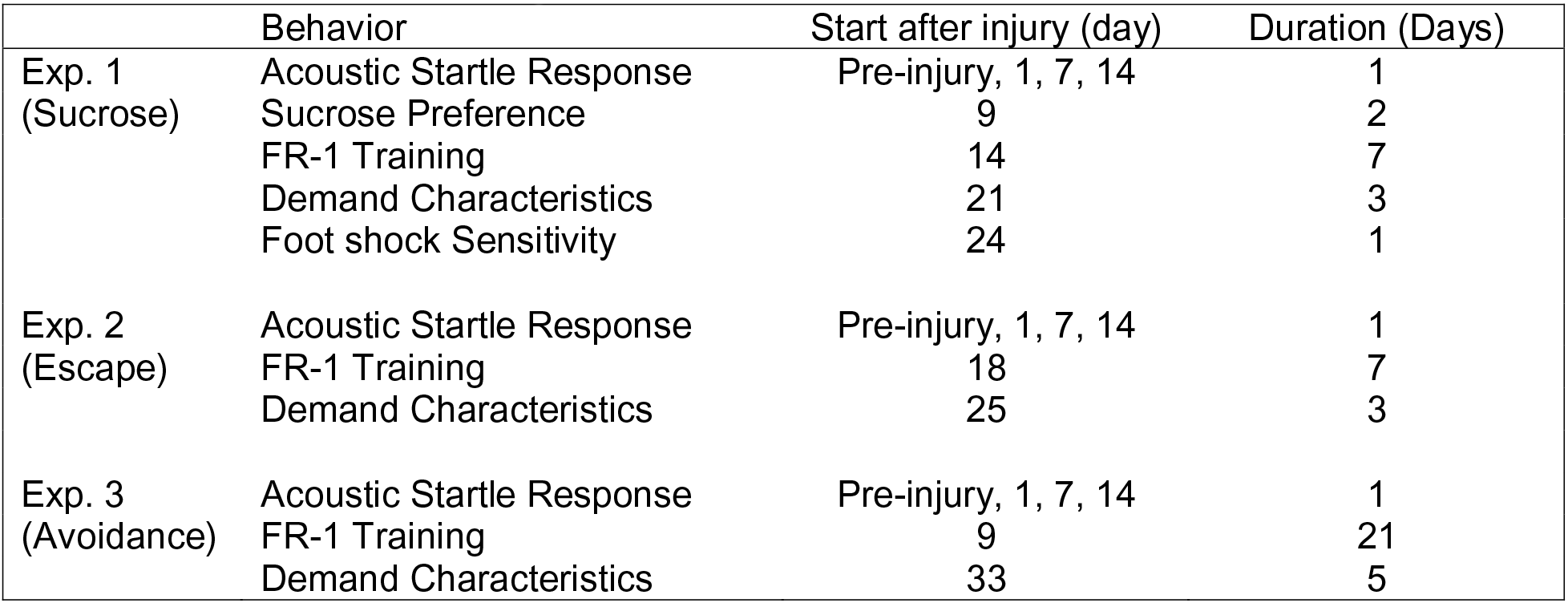
Timeline of Behavioral Testing

#### 2.5.1. Experiment 1: Behavioral Economics of Positive Reinforcement

##### 2.5.1.1. Sucrose Preference

Six sham and 5 mTBI rats were measured for their preference for sucrose using procedures described previously [15]. Rats were presented with two identical water bottles in their home cage. One bottle was filled with tap water and the other filled with 1% sucrose in tap water. Rats were acclimated to the bottles for the first day, and fluid intake was measured during the second day for 24 hours. The position of the water bottles was exchanged after 12 hours during the measurement period. Preference was calculated by the following equation: (volume sucrose consumed – volume water consumed)/(volume sucrose consumed + volume water consumed) × 100.

##### 2.5.1.2. Initial FR-1 training

After assessment of sucrose preference, rats were trained to lever press for sucrose pellets using procedures and equipment described previously [35]. Rats were trained in an operant box that contained a lever (10.5 cm above the floor) on one wall and a house light and speaker (26 cm above the floor) on the opposite wall. A food trough was located below the lever. The operant box was housed in a sound attenuating chamber (Coulbourn). Initial training occurred in 5 daily sessions using a fixed ratio 1 (FR-1) schedule. The presentation of an auditory stimulus (1 kHz, 75 dB) signaled the opportunity to receive reinforcement. A single lever press caused the delivery of a sucrose pellet (45 mg) and then a 20 second intertrial interval (ITI) that was signaled by a flashing light (5 Hz). All stimuli and food dispensers were controlled by Graphic State Notation (Version 4, Coulbourn Instruments). A session consisted of 24 completed trials or 90 minutes, whichever occurred first. Time to criterion (24 reinforced trials) was used to assess learning. Rats unable to obtain 24 reinforced trials by the end of the session were given a score of 90 minutes. By the end of training, mean completion time for a session was 13.8 ± .95 and 16.4 ± 2.38 minutes for Sham and mTBI, respectively. One mTBI rat was unable to complete the 24 trials within 30 minutes so its data were not included in any of the analyses, leaving 6 Sham and 5 mTBI rats.

##### 2.5.1.3. Modified Progressive Ratio Testing

After FR-1 training, rats were tested in a modified progressive ratio procedure, in which positive reinforcement was delivered after increasing numbers of lever presses. The requirement for reinforcement was increased every 6 trials in the following manner: 1, 2, 3, 5, 10, 18 and 32 presses. The lever press requirement increased incrementally regardless of performance. Each rat was tested on 3 daily sessions of the modified progressive ratio procedure to obtain a demand curve function.

Behavioral economics theory is based on the idea that consumption decreases as the price for goods increases. Here, the nature of the relationship between consumption and price is defined by the demand curve. Consumption is the amount of sucrose pellets obtained, expressed as the probability of reinforcement for each trial for a given price. Price is defined as the amount of work required to obtain goods, expressed as the number of lever presses required to receive a sucrose pellet. The exponentiated demand equation was used in this behavioral economic analysis because it allows the use of consumption values equal to 0 [35, 45]. The exponentiated demand equation is

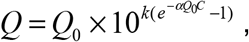

where Q is a measure of consumption of reinforcement, C is the amount of work required to receive reinforcement, and k is a constant that represents the range of consumption and is shared across all subjects. Q_0_ and α are determined from the best fit of the exponentiated demand equation to the composite demand curve. Q_0_ is the consumption of a reinforcer when no effort is required to obtain the reinforcement. While this is a theoretical construct, it is useful as a measure of the hedonic value of a reinforcer. α is a measure of demand elasticity, which describes how rapidly consumption decreases as price increases, and serves as an inverse measure of motivation for obtaining reinforcement. Behavioral economics principles, as well as Q_0_ and α, have been widely used to describe positive reinforcement [17, 20, 21, 46].

##### 2.5.1.4. Foot Shock Sensitivity Testing

In order to provide information for Experiments 2 and 3, rats in Experiment 1 were assessed for sensitivity to foot shock after Modified Progressive Ratio testing with sucrose. Foot shock sensitivity was assessed in an operant box with grid floors capable of delivering scrambled foot shock (Coulbourn Instruments, Whitehall, PA), as described previously [36]. Vocalization and flinch to incrementing shock intensities (0.1 mA steps, 0.1 – 1.0 mA range, 0.5 s duration, 10 s between shocks) were scored by two observers. Each shock intensity was repeated twice. Vocalization was recorded when the animal made an audible vocalization upon shock delivery. A flinch was recorded when the animal visibly tensed its limb muscles immediately upon shock delivery. Threshold was determined to be the lowest shock intensity at which both observers independently agreed on the occurrence of a vocalization or body flinch.

#### 2.5.2. Experiment 2: Behavioral Economics of Negative Reinforcement – Escape from Foot Shock

##### 2.5.2.1. Initial FR-1 training

A separate group of rats (n = 7 sham and 7 mTBI) were trained to lever press in order to escape foot shock. Training occurred as described previously [35]. The operant box was similar to that used in Experiment 1, except without a food trough and with a floor grid that was capable of delivering scrambled foot shock. Rats were trained to press a lever to escape foot shock in 5 daily sessions using an FR-1 schedule. Trials started with foot shock (1.0 mA, 0.5 s duration, 3.5 s intershock interval) paired with a 1 kHz tone (75 dB, continuous). A single lever press immediately terminated the foot shock and auditory stimulus, and initiated a 180 s ITI signaled by a flashing light (5 Hz). A maximum of 20 foot shocks were allowed per trial, after which foot shock and auditory stimulus was terminated, and the ITI initiated. Each session consisted of 25 trials. None of the rats failed to respond on more than 10% of the trials by the end of training.

##### 2.6.2.2. Modified Progressive Ratio Testing

Similar to Experiment 1, a modified progressive ratio procedure was used to construct a demand curve for escape foot shock. The number of lever presses required to terminate foot shock incremented every 6 trials in the following manner: 1, 2, 3, 5, 10, 18 and 32. Each rat was tested on 3 daily sessions of modified progressive ratio testing.

#### 2.5.3. Experiment 3: Behavioral Economics of Negative Reinforcement – Avoidance of Foot Shock

##### 2.5.3.1. Initial FR-1 training

A separate group of rats (n = 7 sham and 8 mTBI) was trained on a lever press avoidance procedure, as described previously [36]. Training occurred in the same operant boxes as in Experiment 2. Each trial was initiated by the presentation of an auditory stimulus (1 kHz, 75dB tone) without foot shock. Foot shocks (1.0 mA, 0.5 s duration, 1 shock/3.5 s) commenced 72 s after the onset of the auditory stimulus. A single lever press after the start of the trial and prior to the initiation of foot shock was recorded as an avoidance response; the avoidance response terminated the auditory stimulus and initiated a 180 s ITI. The ITI was signaled by a flashing light (5 Hz). If an avoidance response was not made, the auditory stimulus continued into the shock period. A single lever press after the start of foot shock terminated shock and the auditory stimulus, and initiated the ITI. Responses during the shock period were recorded as escape responses. If the lever was not pressed after 20 footshocks (72 s), the auditory stimulus and shock were terminated and the ITI commenced. This lack of response was considered a failure. The flashing light was present during the ITI regardless of whether the animal avoided, escaped or failed to respond on the trial. Each training session consisted of 25 trials. Rats were trained for 10 sessions, and sessions occurred 3 times per week with a minimum of 48 hours between sessions, as in previous studies [47].

##### 2.5.3.2. Modified Progressive Ratio Testing

A modified progressive ratio procedure was used to construct a demand curve for avoiding foot shock, similar to that described for escape from foot shock. Rats were tested in the same operant boxes as during FR-1 training. The procedure was identical to that described for the FR-1 avoidance training, except the number of lever presses to avoid foot shock was increased in the following manner: 1, 2, 3, 5, 10, 18 and 32. The number of lever presses incremented every 6 trials regardless of performance. Throughout the testing session, a single lever press was necessary to terminate foot shock during the shock period (escape response). Each rat was tested on 3 sessions of the modified progressive ratio schedule with 48 hours between sessions.

### 2.6. Data Analysis

#### 2.6.1. Statistical Analysis

Statistical analyses of ASR, sucrose preference, foot shock sensitivity, acute injury measures and FR-1 training were performed using SPSS for Windows (v 12.0.1). ASR sensitivity and magnitude were analyzed in a 2 × 4 × 3 mixed design ANOVA, with between-subjects factor of injury (2 levels: SHAM or mTBI) and within-subject factors of day of measurement (4 levels: pre-injury, 1, 7 and 14 days after injury) and stimulus intensity (82, 92, 102dB). Data from FR-1 training were assessed using a 2 × 5 (or 10) mixed design ANOVA, with between subjects factor of injury (2 levels: SHAM or mTBI) and a within subject factor of session (5 levels for Experiments 1 and 2 and 10 levels for Experiment 3). Because these studies focused on the effect of mTBI, only main effects and interactions involving the injury factor are reported. Two-tailed independent samples t-tests were conducted for sucrose preference, foot shock sensitivity, and acute measures of fluid percussion impact. Statistical significance was reached when *p* < .05.

#### 2.6.2. Behavioral Economics

Each rat was tested in the modified progressive ratio procedure for 3 sessions. Data from the three sessions were combined to create a demand curve for each rat. A composite demand curve was then constructed from all rats for each group. The exponentiated demand equation was fit to the composite demand curve using a modified GraphPad Prism (v. 6.0 h) template from the Institutes for Behavior Resources. The measures of hedonic value Q_0_ and elasticity α for sham and mTBI groups were obtained from the best fit of the exponentiated demand equation to the composite demand curve for each group. All fits of the equation to actual data were excellent (r^2^ between 0.84 and 0.99). Statistical interpretations of Q_0_ and α were obtained from the Extra Sum of Squares F-test (GraphPad Prism, v. 6.0h). Statistical significance was reached when *p* < .05.

## 3. Results

A timeline of the behavioral procedures for the three experiments is presented in Table 2.

### 3.1. Experiment 1: Behavioral Economics of Positive Reinforcement

#### 3.1.1. Injury Severity and Acute Signs

As measures of injury severity, impact pressures and acute signs of injury were recorded and are summarized in Table 1. Time to recover righting reflex was significantly longer for mTBI compared to sham (t(9) = −7.9, p < .001).

#### 3.1.2. Acoustic Startle Response

We have previously reported a long-lasting suppression of ASR following mTBI after lateral fluid percussion injury [37–39]. In the present study, we used ASR suppression as an additional measure of injury. ASR sensitivity and magnitude were evaluated up to 14 days following injury. Similar to our previous studies, ASR sensitivity was suppressed following mTBI, as supported by a main effect of injury (F(1,9) = 22.7, p = .001), an injury × day interaction (F(3,27) = 5.3, p = .005) and an injury × stimulus intensity interaction (F(2,18) = 7.5, p = .004) (Figure S1). The triple interaction of injury × day × stimulus intensity trended toward significance (F(6,54) = 2.0, p = .08.

Similar to ASR sensitivity, magnitude of ASR was suppressed following mTBI (Figure 1, top). The significant interaction of injury × day (F(3,27) = 3.8, p = .021) supported the adverse effect of mTBI. The main effect of injury (F(1,9) = 4.2, p = .07) trended toward significance. All other interactions involving injury were not significant. Thus, suppression of both ASR sensitivity and magnitude resulted from lateral fluid percussion injury.

**Figure 1.**
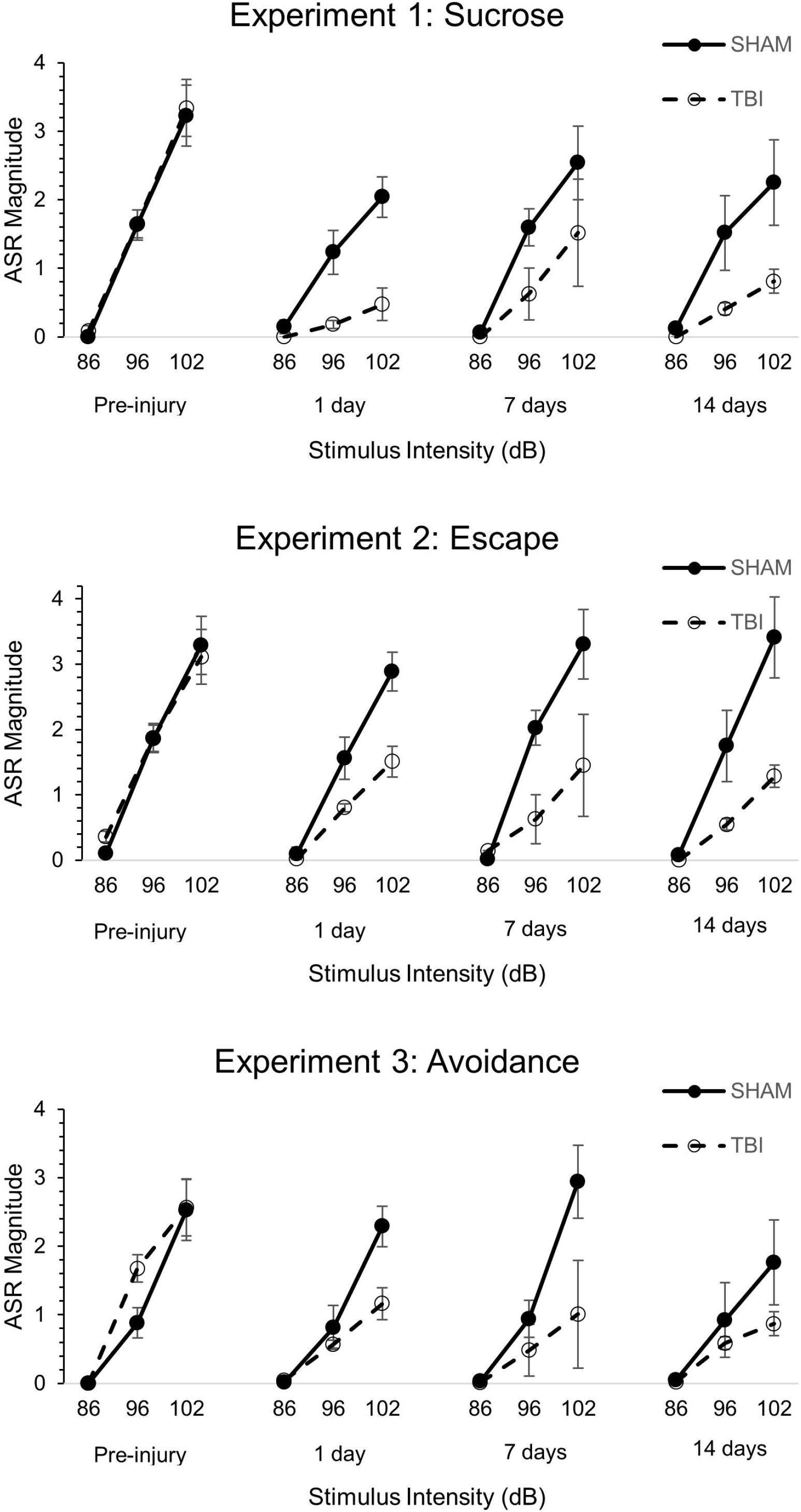
Magnitude of acoustic startle responses (ASR) was suppressed following mTBI. ASR was obtained from rats in Experiments 1, 2 and 3 prior to injury and 1, 7 and 14 days after mTBI. The magnitude of ASR was suppressed in all experiments, as demonstrated by significant main effects of injury and interactions involving injury with stimulus intensity and day (see text discussion of statistical results). Similar effects of mTBI were observed for sensitivity of ASR (Figure S1).

#### 3.1.3. Sucrose Preference

Sucrose preference was assessed 9 days after injury. Both Sham and mTBI rats preferred the sucrose solution, and the groups did not differ in their preference (t(9) = 1.2, p = .261) or total amount of liquid consumed (t(9) = −1.0, p = .336) (Table 3).

**Table 3.** Experiment 1 – Sucrose Preference and Foot Shock Sensitivity. Mean ± S.E.M.

| Condition | Sucrose Preference (%) | Vocalization Threshold (mA) | Flinch Threshold (mA) |
| --- | --- | --- | --- |
| Sham | 93.5 ± 1.6 | .30 ± .04 | .52 ± .03 |
| mTBI | 87.2 ± 5.5 | .22 ± .02 | .50 ± .08 |

#### 3.1.4. FR-1 Training

Rats were trained to lever press for sucrose pellets in 5 sessions using an FR-1 reinforcement schedule. Learning was assessed by the time to reach a criterion of 24 reinforced trials (Figure S2). Sham and mTBI groups decreased time to complete the session with training (main effect of session, F(4,36) = 18.6, p < .001), and learning was not different between groups, as the main effect of injury and injury × session interaction were not significant (ps > 0.05).

#### 3.1.5. Demand characteristics for Positive Reinforcement – Sucrose

Effects of mTBI on the reinforcing properties of positive reinforcement were evaluated in a modified progressive ratio schedule for sucrose pellets. Demand characteristics were obtained for Sham and mTBI rats (Figure 2A). Q_0_, a measure of the hedonic value of sucrose, was not different between Sham and mTBI rats (F(1,10) = 0.82, p = .385) (Figure 2B). α, a measure of elasticity that is inversely related to motivation, was also not different between Sham and mTBI groups (F(1,10) = 0.43, p = .527) (Figure 2C). Fit of the exponentiated equation to actual data was excellent (r^2^ for sham = 0.93 and mTBI = 0.96).

**Figure 2.**
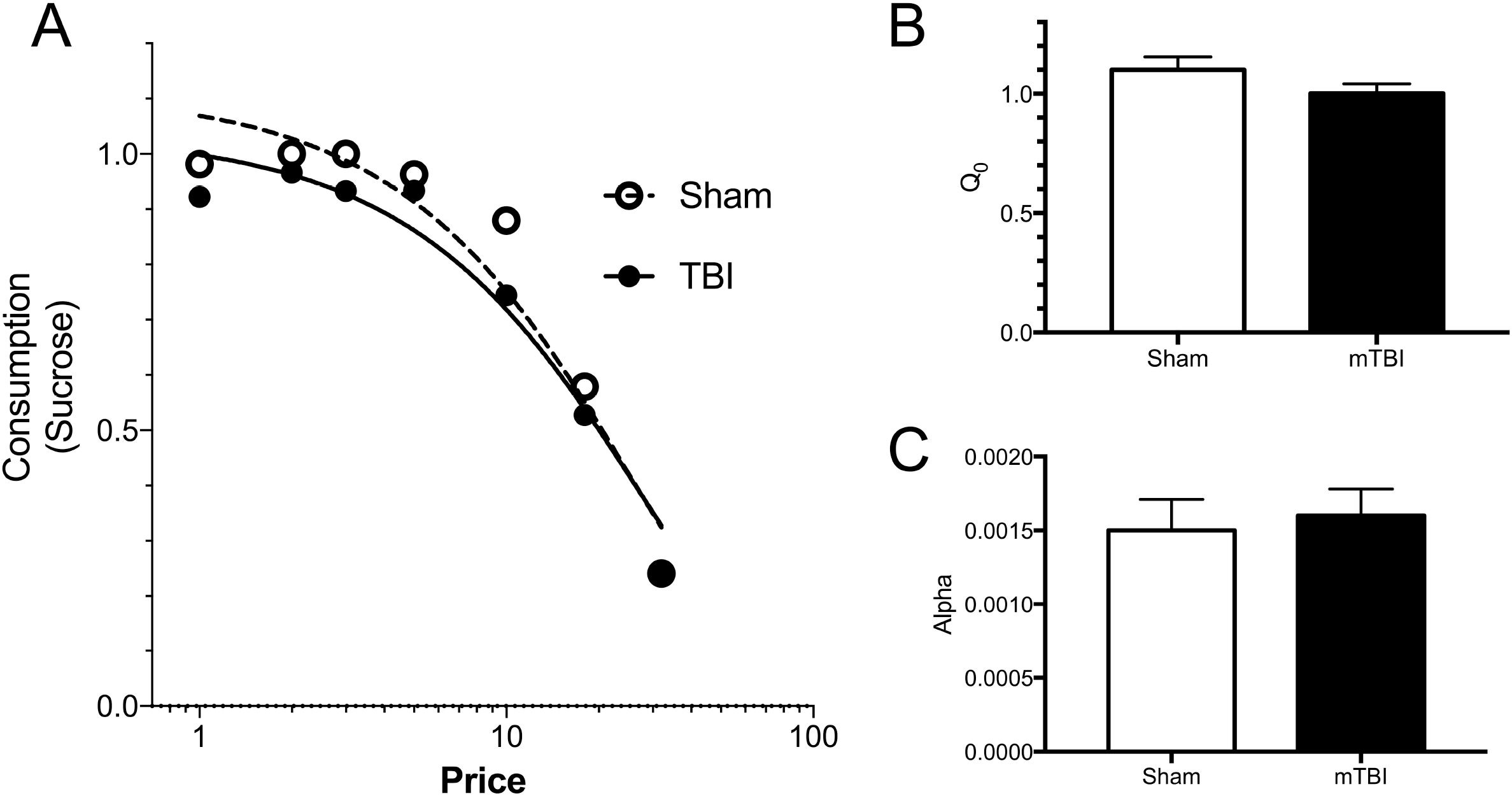
Effects of mTBI on demand for sucrose. After learning to lever press to obtain sucrose pellets, a demand curve was obtain using a modified progressive ratio procedure (A). Neither Q_0_ (B) nor α (C) were different between sham and mTBI rats, demonstrating that neither hedonic value of sucrose nor the motivation to obtain sucrose was altered by mTBI.

#### 3.1.6. Sensitivity to Foot Shock

As a prelude to Experiments 2 and 3, rats from Experiment 1 were evaluated for their sensitivity to foot shock by assessing threshold to flinch and vocalize in response to varying levels of foot shock. Mean foot shock threshold values are presented in Table 3. Sham and mTBI groups did not differ in the threshold to vocalize (t(9) = 1.8, p = .104) or flinch (t(9) = .201, p = .845). Thus, the results support the view that mTBI did not change the sensitivity to foot shock.

In summary, the results of Experiment 1 demonstrated that mTBI caused a suppression of ASR, but did not alter reinforcement value or motivation to obtain positive (sucrose) reinforcement. Additionally, mTBI did not modify their preference for drinking sucrose water or sensitivity to foot shock.

### 3.2. Experiment 2: Behavioral Economics of Negative Reinforcement – Escape from Foot Shock

#### 3.2.1. Injury severity and Acute Signs

Impact pressures and acute signs for Experiment 2 are summarized in Table 1. Latency to recover the righting reflex was significantly longer for mTBI compared to sham (t(12) = −4.62, p = .001).

#### 3.2.2. Acoustic Startle Response

ASR sensitivity for rats in Experiment 2 was not suppressed following mTBI, as the main effect of injury and all interactions involving injury were not significantly different between sham and TBI groups (Figure S1). In contrast, ASR magnitude was suppressed by mTBI, as demonstrated by a significant main effect of injury (F(1,13) = 7.7, p = .016) and interactions of injury × day (F(3,39) = 4.0, p = .015), injury × stimulus intensity (F(2,26) = 8.4, p = .002) and injury × day × stimulus intensity (F(6,78) = 2.3, p = .04) (Figure 1 middle). Thus, mTBI significantly suppressed ASR, although the effect was only observed for the magnitude of ASR. This was not surprising since we have previously observed that the effects of mTBI on ASR are greater for magnitude than for sensitivity [37, 38].

#### 3.2.3. FR-1 Training

Sham and mTBI rats were trained to lever press to escape foot shock using an FR-1 reinforcement schedule. Learning performance was assessed by calculating the proportion of trials with an escape response (Figure S3). Both groups learned the escape response with training (main effect of session, F(4,52) = 15.3, p < .001), but acquisition of escape was not different between Sham and mTBI groups, as the main effect of injury and the injury × session interaction were not significantly different.

#### 3.2.4. Demand Characteristics of Negative Reinforcement – Escape from Foot Shock

The effects of mTBI on the reinforcing properties of escape from foot shock was investigated using a modified progressive ratio procedure. Q_0_ was not different between Sham and mTBI rats (F(1,10) = 0.005, p = .978) (Figure 3B). α for escaping foot shock was also not different between groups (F(1,10) = 0.1, p = .754) (Figure 3C). The fit of the exponentiated equation to the actual data was very good (r^2^ for sham = 0.93 and mTBI = 0.84) (Figure 3A). Thus, mTBI did not alter the hedonic value or motivation of escaping from foot shock, evidence that reactive control of stress was not altered by mTBI.

**Figure 3.**
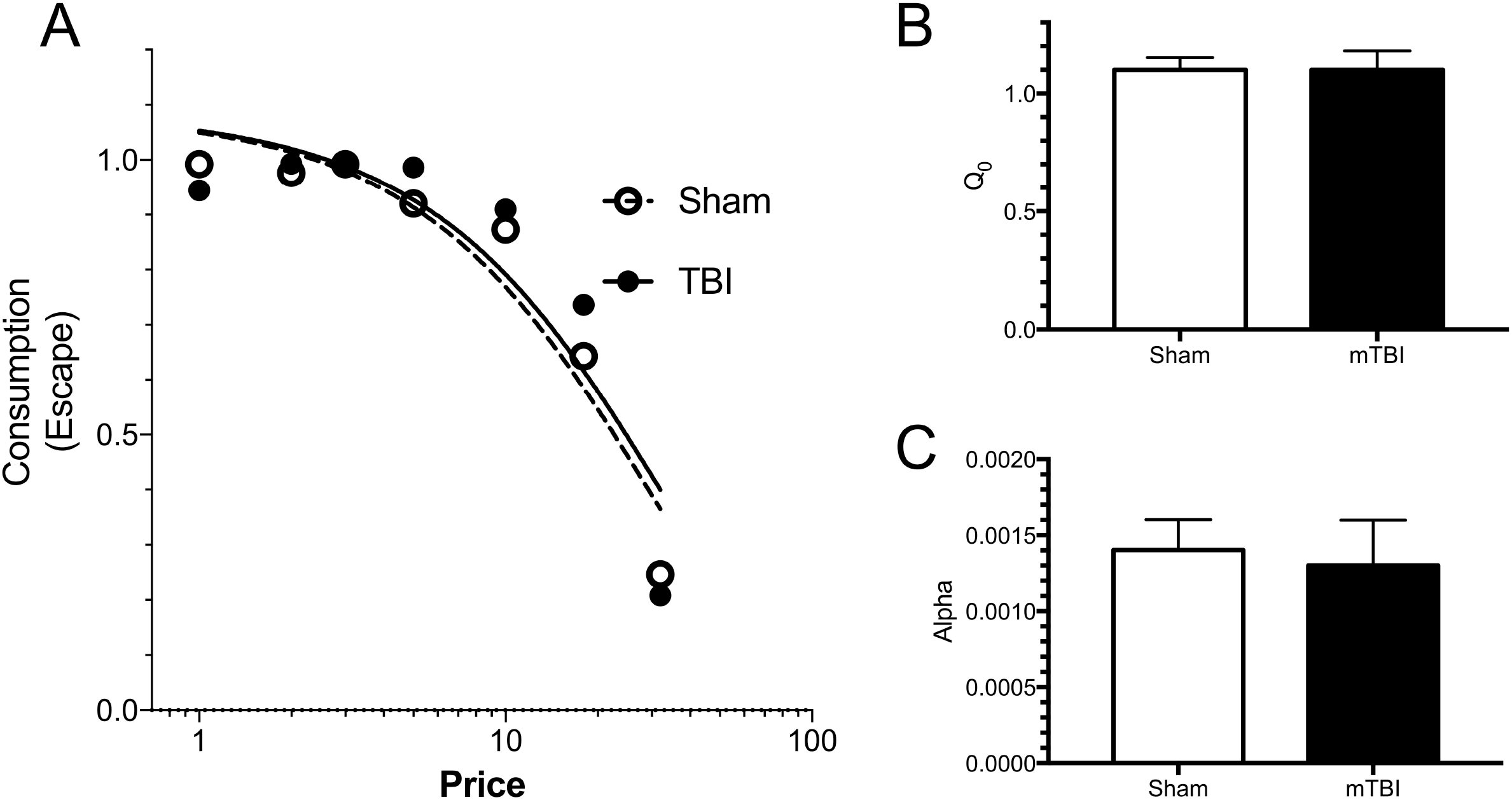
Effects of mTBI on escape from foot shock. Rats learned to lever press to escape foot shock (1mA, 0.5s). After acquisition, a demand curve was obtained using a modified progressive ratio procedure (A). Neither Q_0_ (B) nor α (C) were different between sham and mTBI rats, demonstrating that mTBI did not alter the hedonic value of escaping foot shock or the motivation to escape foot shock. These results suggest that mTBI does not alter reactive control of stress.

### 3.3. Experiment 3: Behavioral Economics of Negative Reinforcement – Avoidance of Foot Shock

#### 3.3.1. Injury severity and Acute Signs

The impact pressure experienced and acute signs shown by rats in Experiment 3 are summarized in Table 1. Latency to recover the righting reflex was significantly longer for mTBI rats compared to sham rats (t(13) = −12.7, p < .001).

#### 3.3.2. Acoustic Startle Response

ASR sensitivity for rats in Experiment 3 was suppressed by mTBI (Figure S1). The main effect of injury (F(1,13) = 6.3, p = .026) and the interaction of injury × stimulus intensity (F(2,26) = 4.2, p = .026) were significantly different between Sham and mTBI groups.

ASR magnitude was also significantly suppressed following mTBI (Figure 1 bottom). The interactions of injury × day (F(3,39) = 5.3, p = .004), injury × stimulus intensity (F(2,26) = 7.5, p = .003) and injury × day × stimulus intensity (F(6,78) = 3.3, p = .006) were significantly different, even though the main effect of injury was not significant. Therefore, mTBI caused the suppression of ASR sensitivity and magnitude in Experiment 3.

#### 3.3.3. FR-1 Training

Sham and mTBI rats were trained to avoid foot shock using an FR-1 schedule. FR-1 training was conducted for 10 sessions, as avoidance learning requires more training than the escape response. Performance was measured by the proportion of trials with an avoidance response (Figure S4). Sham and mTBI rats learned the avoidance response (main effect of session, F(9,108) = 13.9, p < .001). Both groups did not differ in their acquisition of avoidance, as neither the main effect of injury nor the injury × session interaction was significant.

#### 3.3.4. Demand Characteristics of Negative Reinforcement – Avoidance of Foot Shock

The effect of mTBI on the negative reinforcement of avoiding foot shock was investigated using a modified progressive ratio procedure, and the demand curve is shown in Figure 4 A. The hedonic value Q_0_ for avoiding foot shock was significantly reduced for mTBI rats as compared to Sham rats (F(1,10) = 20.0, p = .0012) (Figure 4B). In contrast, α for avoiding foot shock was not significantly different between injury groups (F(1,10) = 1.8, p = .209) (Figure 4C). The fit of the exponentiated equation to actual data were excellent (r^2^ for sham = 0.97 and mTBI = 0.99). Thus, unlike the lack of effects of mTBI on the demand characteristics for obtaining sucrose and escaping from foot shock, mTBI reduced the reinforcing value of avoiding foot shock.

**Figure 4.**
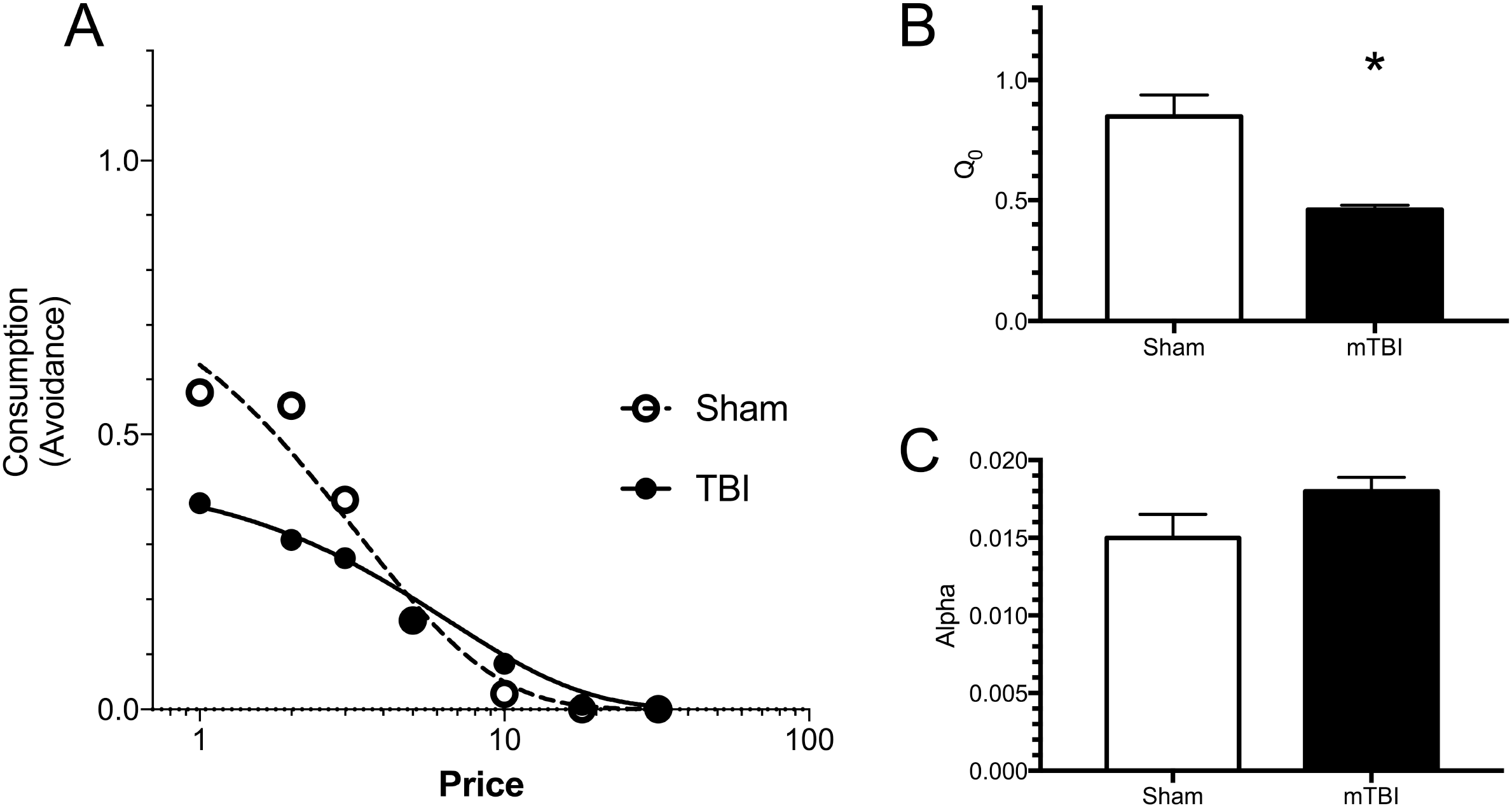
Effects of mTBI on avoidance of foot shock. After learning to lever press to avoid foot shock (1mA, 0.5s), a demand curve for avoidance was obtained using a modified progressive ratio procedure (A). Q_0_ was reduced by mTBI, suggesting a reduction of the hedonic value of an avoidance response. In contrast, α was not altered by mTBI. The differential effect of mTBI on avoidance but not escape of foot shock supports the idea that mTBI impairs proactive, but not reactive, control of stress by reducing the reinforcing value of avoidance responding.

## 4. Discussion

In the last 20 years, the concept of reinforcement has been refined to include different components, such as hedonic, motivational and learning [48, 49]. Similar distinctions of reinforcement have been applied to psychopathology where consummatory, motivational and decisional components of reinforcement may be differentially affected in individuals to account for different symptoms [7, 50]. By exploring the characteristics of demand and using behavioral economic approaches, the current study sought to understand the effects of mTBI on hedonic and motivational components of reinforcement.

Mood disorders including depression and anxiety can differentially alter views of positively- and negatively-valenced stimuli or events. Stress due to encountering aversive events can precipitate or exacerbate depression and anxiety [30, 31]. Cognitive control can help an individual deal with or proactively reduce the effects of stressors [51]. Conversely, impaired cognitive control may lead to preoccupation of negative events, often referred to as negative bias in which attention to aversive events or thoughts is greater than to positive events or thoughts [52, 53]. Thus, a compelling case can be made that the study of negative reinforcement (i.e., termination or avoidance of negatively-valenced events) is as relevant and important to the study of mood disorders as the study of positive reinforcement (occurrence of positively-valenced events).

A novel rat model of despair/depression utilized extinction of positive or negative reinforcement behaviors [54–56]. Withdrawal of reinforcement during the extinction phase provides uncontrollable stress and aversiveness that can lead to helplessness. Interestingly, rats bred to express helplessness in a negative reinforcement condition were less motivated to obtain flavored pellets, a positive reinforcer, when compared to the non-helpless variant and wild type control rat [57]. Relevant to the present study, the selectively bred helpless and non-helpless rats did not differ on consumption of the flavored food when easily available, a measure of hedonic value. Thus, the selective breeding of helpless behavior did not result in anhedonia for positive reinforcement.

In the present study, the effects of mTBI on the hedonic and motivational components of positive and negative reinforcement were investigated. Two types of negative reinforcement were assessed to determine the type of cognitive control affected by mTBI: escape (reactive control) and avoidance (proactive control) of foot shock (stressor). mTBI was produced using a lateral fluid percussion injury model. The impact pressure and acute signs were similar to our previous studies and to those previously reported for mild severity TBI [37, 38, 58–61]. Suppression of ASR was also observed after mTBI, as reported previously [37–39, 62].

With regard to the main focus of this study, mTBI reduced the hedonic value of avoiding foot shock, a form of anhedonia for negative reinforcement. In contrast, mTBI did not alter the hedonic value or the motivation to escape foot shock, as neither Q_0_ nor α were significantly different between sham and mTBI groups. The fact that the hedonic value of *avoiding* but not *escaping* foot shock was impacted by mTBI point to a very specific impairment in negative reinforcement, in particular proactive rather than reactive control of a stressor. Thus, individuals with mTBI would find proactive coping of stress less reinforcing, and therefore be less likely to use this coping strategy to deal with stress [34, 52]. Importantly, cognitive behavioral therapy (CBT) is associated with enhanced activity in the cognitive control brain circuits of patients with major depression, suggesting CBT may be particularly efficacious in TBI-induced depression [63].

The effect of mTBI on avoidance is unlikely to be due to changes in sensitivity to foot shock after TBI. mTBI did not change the hedonic value or the motivation to escape foot shock nor did mTBI alter thresholds to vocalize and flinch to foot shock. Moreover, the lack of effects of mTBI on escape from foot shock helps to rule out other explanations such as motor impairment or general learning deficit.

At present, we can only speculate on the mechanisms by which mTBI impaired the hedonic value of avoidance and proactive control of stress. Expectancy is critical for proactive control and the dorsolateral prefrontal cortex (DLPFC) is an involved in expectancy of goal-relevant emotional information. Sustained activation of the DLPFC in humans occurred during high expectancy conditions [64]. Another possibility is the dopaminergic system, which is important for conditioned avoidance. Lesions of dopaminergic terminals and dopaminergic antagonists impair foot shock avoidance, and dopamine release is correlated with avoidance responses [65–69]. TBI modifies neural circuits involving both DLPFC and dopamine systems [70–73]. While dopaminergic systems have largely been implicated in both positive and negative reinforcement, any mechanistic explanation of our results will have to include differential actions of TBI on avoidance of foot shock, but not escape or positive reinforcement [74, 75].

Our results showing a lack of effect of mTBI on positive reinforcement are consistent with one study, but at odds with others. Severe lateral fluid percussion TBI in Wistar rats led to no changes in sucrose preference at 1 to 6 months after injury, but fluid percussion TBI interacted with lipopolysaccharide (LPS) administration to decrease sucrose preference to a greater degree than LPS in sham mice 30 days after injury [14, 15]. Controlled cortical impact (CCI) injury in Sprague Dawley rats resulted in lower sucrose preference at 1 week and 5.5 months after injury [12, 76]. Finally, repetitive closed head injury (2 injuries, 24 hours apart) decreased sucrose preference 2 weeks after injury [77]. In general, reduced sucrose preference was observed after TBI in rodents suggesting anhedonia for positive reinforcement, in line with anhedonia observed in TBI patients. In the present study, sucrose preference after mTBI was not changed nor was hedonic value or motivation to obtain sucrose using an operant procedure with behavioral economics analyses. It is possible that the differences between our results and previous findings are due to the nature of the injury (fluid percussion vs CCI vs closed head injury), severity or location of injury.

## 5. Conclusion

The present study investigated the effects of mTBI on different components of positive and negative reinforcement. mTBI reduced the hedonic value of avoiding foot shock, but did not modify other aspects of avoidance or escape of foot shock or obtaining sucrose. Thus, results of the present study suggest that mTBI impairs proactive coping of stress through anhedonia for negative reinforcement, which may ultimately lead to depression.

## Acknowledgement

Supported by Merit Award (I01BX000132) to KCHP and Career Development Award (IO1BX007080) to AMF, both from the Biomedical Laboratory Research & Development Service of the VA Office of Research and Development, and Graduate/Post-Doctoral Fellowship grant (CBIR14FELO14) to PA from New Jersey Commission on Brain Injury Research.

The views expressed in this article are those of the authors and do not necessarily reflect the position or policy of the Department of Veterans Affairs or the United States government.

