## Supplemental Figures for "Anhedonia following mild traumatic brain injury in rats: A behavioral economic analysis of positive and negative reinforcement"

| 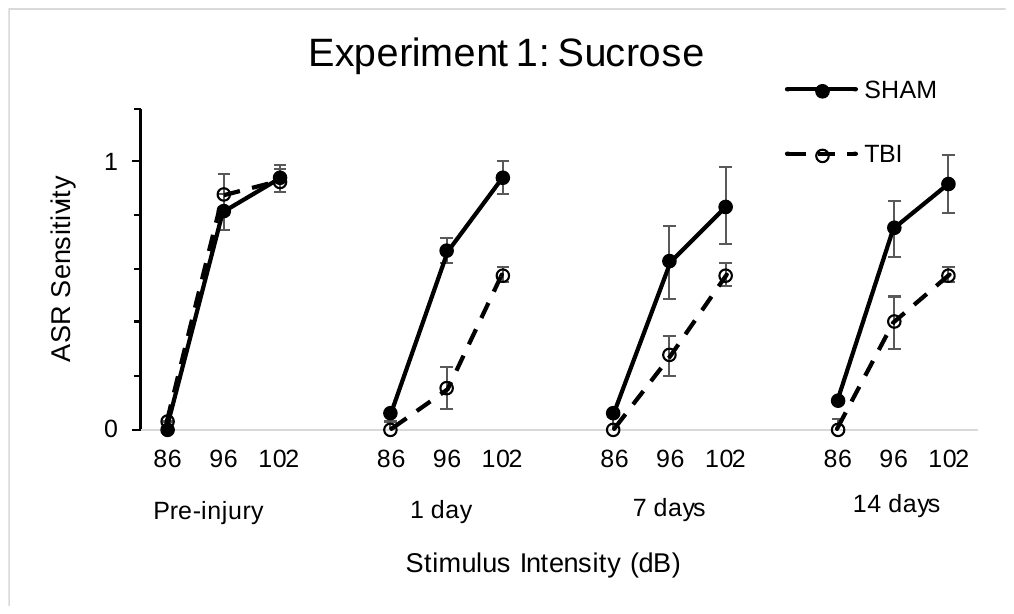 | Figure S1. Sensitivity of acoustic startle responses (ASR) was suppressed following mTBI. ASR was obtained from rats in Experiments 1, 2 and 3 prior to injury and 1, 7 and 14 days after mTBI. (Top) ASR sensitivity was suppressed after mTBI in Exp 1, as demonstrated by main effect of injury (F(1,9)=22.7, p = .001), injury X day interaction (F(3,27)=5.3, p= .005) and injury X stimulus intensity interaction (F(2,18)=7.5, p=.004). The injury X day X stimulus intensity interaction was trending, F(6,54)=2.0, p=.08. (Middle) ASR sensitivity was not altered by mTBI in Exp 2. (Bottom) ASR sensitivity was suppressed after mTBI in Exp 3, as supported by a main effect of injury (F(1,13)=6.3, p=.026) and an injury X stimulus intensity interaction (F(2,26)=4.2, p=.026). |
| --- | --- |
| 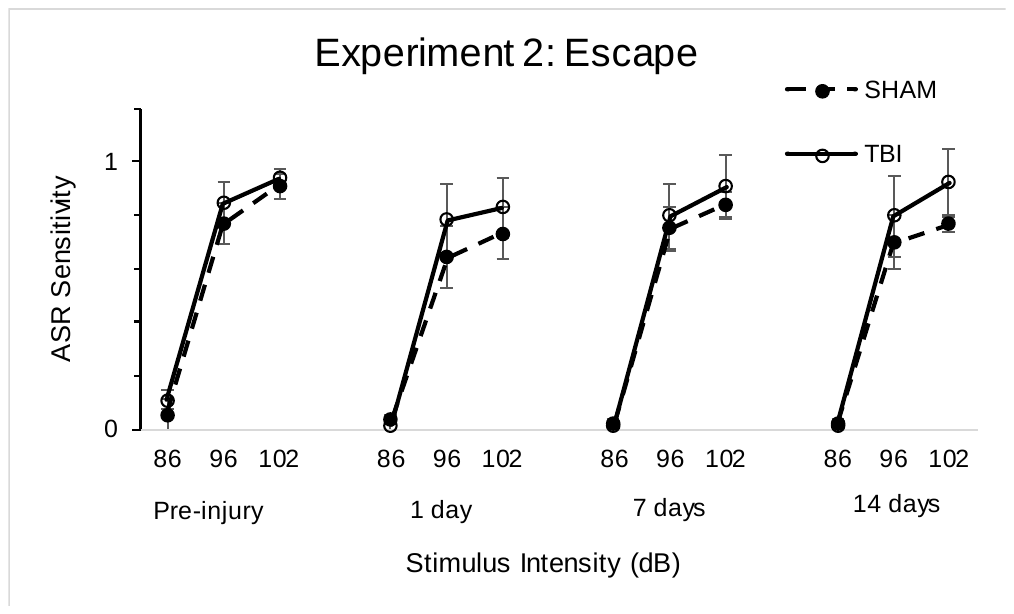 |  |
| 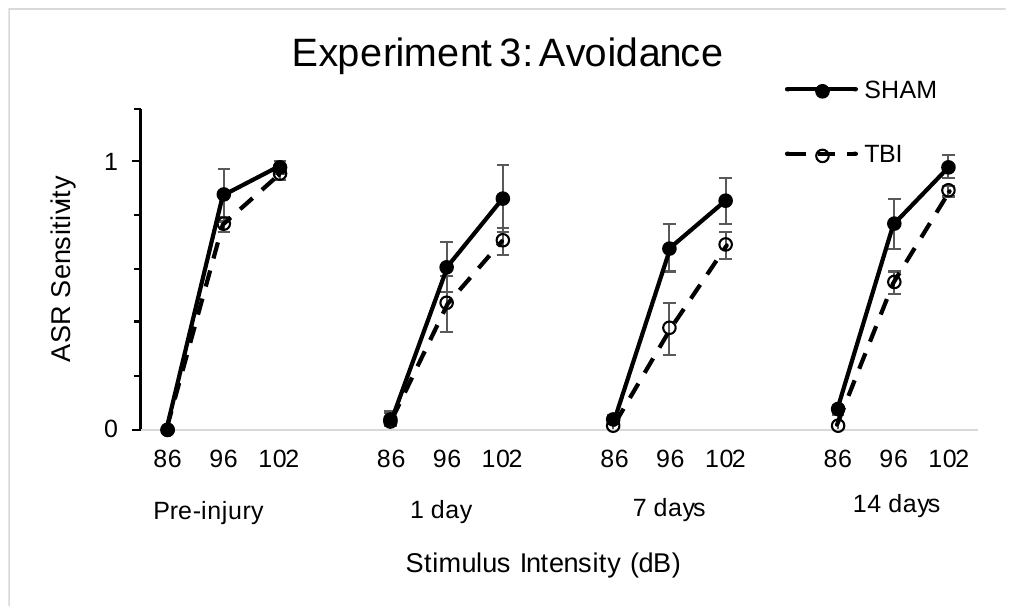 |  |

| 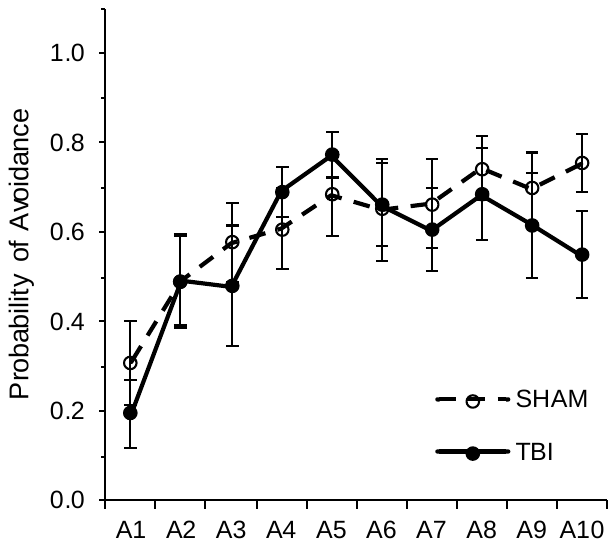 | Figure S4. mTBI did not alter avoidance learning. Rats were trained to lever press to avoid foot shock (1 mA, 0.5s every 3.5 s). A single lever press during the initial 72 s of the trial prevented foot shock (see text for more details). Each session consisted of 25 trials. Both sham and mTBI rats learned to avoid foot shock, as demonstrated by an increase in the probability of trials with an avoidance response (main effect of days, F(9,108)=13.86, p < .001). Sham and mTBI groups did not differ in avoidance learning (main effect of injury, F(1,12)=0.02; injury X day interaction, F(9,108)=1.07). |
| --- | --- |

| 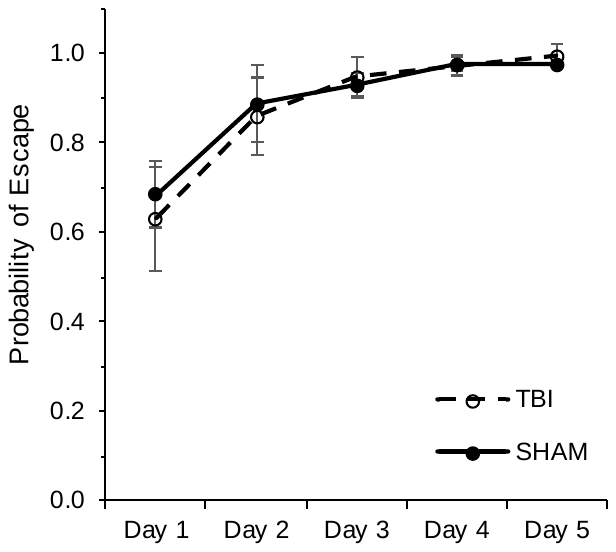 | Figure S3. mTBI did not alter learning to escape foot shock. Rats were trained to lever press for terminate foot shock (1 mA, 0.5s every 3.5 s). A single lever press caused the cessation of foot shock (see text for more details). Each session consisted of 25 trials. Both sham and mTBI rats learned to lever press to escape foot shock, as demonstrated by an increase in the probability of trials with an escape response (main effect of days, F(4,52)=15.3, p < .001). Sham and mTBI groups did not differ in learning to escape foot shock (main effect of injury, F(1,13)=0.02; injury X day interaction, F(4,52)=0.21). |
| --- | --- |

| 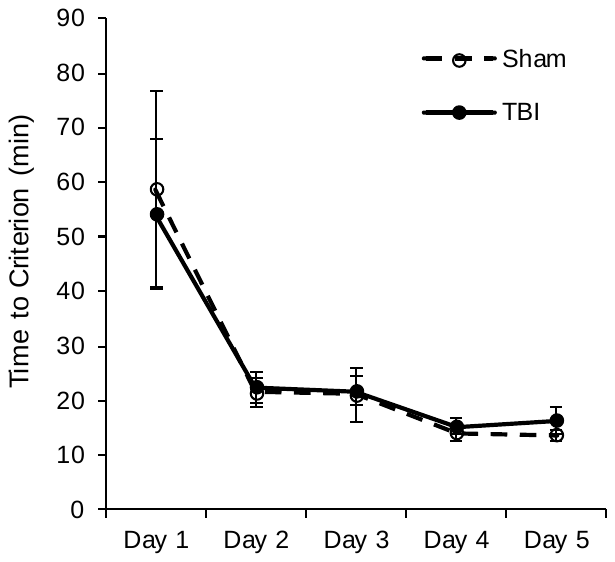 | Figure S2. mTBI did not alter learning to lever press for sucrose. Rats were trained to lever press for sucrose reinforcement. A single lever press delivered a sucrose pellet (see text for more details). Each session terminated after delivery of 24 pellets or 90 minutes. Time to criterion was the session time to acquire 24 pellets or 90 minutes if the rat did not obtain 24 reinforcements. Both sham and mTBI rats learned to lever press for sucrose pellets, as demonstrated by a decrease in criterion time (main effect of day, F(4,36)=18.6, p < .001), but learning did not differ between groups (main effect of injury, F(1,9)=0.17; injury X day interaction, F(4,36)=0.05). |
| --- | --- |
